# Broadly neutralizing human monoclonal antibodies against BK polyomavirus genotypes

**DOI:** 10.1101/2024.12.18.629307

**Authors:** J. Andrew Duty, Thomas Kraus, Madhu Kumar, Nicolo A. Tortorella, Tajudeen O. Jimoh, Emma B. Barrall, Diana V. Pastrana, Christopher B. Buck, Thomas Moran, Domenico Tortorella

## Abstract

BK polyomavirus (BKV) causes polyomavirus-associated nephropathy (BKV-nephropathy) and polyomavirus-associated hemorrhagic cystitis (BKV-HC) following kidney transplantation and allogeneic hematopoietic stem cell transplantation (HSCT). BKV strains consist of four distinct genotypes (BKV-I, - II, -III, and -IV) with more than 80% of individuals seropositive for the BKV-I genotype, and lower prevalences of infection or co-infection with the other genotypes. BKV-nephropathy occurs widely in immunosuppressed transplant recipients, with the recommended treatment including reduction of immunosuppressive drugs. High serum titers of BKV-neutralizing antibodies and treatment with BKV-neutralizing intravenous immunoglobulin (IVIG) are associated with reduced levels of BKV-DNAemia in kidney transplant recipients, suggesting anti-BKV antibodies can limit viral load. Thus, we set out to generate broadly neutralizing human monoclonal antibodies (mAbs) against the viral protein 1 (VP1) major capsid protein of BKV genotypes I-IV using VelocImmune® transgenic mice that encode for human immunoglobulins. Hybridoma clones from VelocImmune® mice immunized with combinations of BKV I-IV VP1 and respective genotypes of BKV pseudoviruses (BK-PsV) were screened using a high-throughput binding assay against BKV-I VP1 expressed in HEK-293 cells. The VP1-binding hybridoma clones were then assessed for neutralization with BK-PsV consisting of respective VP1 proteins from BKV-I, -II, -III, or -IV on the virion surface. Overall, thirty-six broadly cross-neutralizing mAbs against BKV-I, -II, -III, and -IV -were identified. Twenty of the cross-reactive immunoglobulins were subjected to nucleotide sequencing resulting in six clonotype families with fourteen genetically distinct immunoglobulins. Several of the most effective mAbs were fully humanized with the IgG1 Fc domain and broad neutralization of BK-PsV-I, II, III, and IV genotypes with IC50s ranging from ∼7-200 pM. Thus, these cross-neutralizing mAbs represent potential biologics to developed into BKV therapeutics.

## INTRODUCTION

BK polyomavirus (BKV), a non-enveloped DNA virus with a genome that has three functional regions: the early viral gene region (EVGR), which encodes the large and small T antigens; the late viral gene region (LVGR), which encodes capsid proteins and non-structural agnoproteins; and the non-coding control region (NCCR)(1). The structure of the BKV capsid is comprised of viral proteins: VP1, VP2, and VP3. The major capsid protein VP1 organizes into 72 protruding pentamers (360 VP1 molecules) arranged in an icosahedral structure, enabling attachment to target cells (2). VP2 and VP3 form part of the inner surface of the BKV capsid. Cellular attachment and infectious entry are mostly mediated by VP1 interactions with cell-surface glycosaminoglycans and sialylated glycans, such as the headgroups of the ganglioside GT1b (3). Notably, however, the BKV genotype IV utilizes sialylated proteoglycans instead of gangliosides for infectious entry (4, 5). BKV is generally categorized into genotypes I, II, III, and IV based on its VP1 sequence, with BKV-I being the most prevalent (60-80%)(6).

BKV infection occurs through oral or respiratory routes and is nearly ubiquitous throughout the US population, with seropositivity by the age of ten between 75-90% (7). Following primary infection, individuals with intact immunity are largely asymptomatic, yet the virus can persist at low levels in the urinary tract (8). In the case of kidney transplantation, however, immunosuppression enables BKV proliferation, increasing viral load, as demonstrated by viral genome numbers in the blood (DNAemia). In fact, BKV-DNAemia can be observed in up to 30% of transplant recipients with 5–10% of patients developing BKV-nephropathy and to kidney graft dysfunction (9, 10). Based on recent guidelines by the Transplantation Society International BK Polyomavirus Consensus Group (11), BKV-nephropathy is classified into four distinct diagnostic levels: 1) possible BKV-nephropathy is indicated by high urine BKV loads (>10 million copies/mL) or viral presence detected by microscopy; 2) probable BKV-nephropathy is consistent with plasma BKV-DNAemia (>1,000 copies/mL) for over two weeks; 3) presumptive BKV-nephropathy is BKV-DNAemia (>10,000 copies/mL); and 4) biopsy-proven BKV-nephropathy requires histological evidence of viral cytopathic effects, positive immunohistochemistry, and a diagnostic test confirming BKV. High levels of BKV-DNAemia also correlate with an increased risk of bladder cancer (12). Given that >27,000 kidney transplants were performed in the US 2024 US (Organ Procurement & Transplantation Network (https://optn.transplant.hrsa.gov/), there is an urgent medical need for effective therapies against BKV.

Several factors of the kidney donor and recipient predispose transplant recipients to BKV-DNAemia and biopsy-proven BKV-nephropathy (11). The donor-contributing factors include urinary BKV shedding, high donor antibody levels against BKV VP1, and genotype mismatching. The transplant recipient factors are age, sex, seronegative BKV VP1 antibody status, previous kidney transplantation, and lack of protective HLA types. Other recipient-associated factors include the use of tacrolimus or corticosteroid treatment following transplant. Importantly, if the graft recipient lacks antibodies against the viral genotype found in the donor organ (13–15), or there is a mismatch in viral genotype between donor and recipient, there is an increased risk of developing BKV-DNAemia and BKV-nephropathy (16, 17). Numerous studies have found that treating kidney transplant recipients with intravenous immunoglobulin (IVIG) decreased BKV-DNAemia, suggesting the protective role of anti-BKV immunoglobulins against BKV-nephropathy (10, 18). Further, preliminary results from a recent Phase II clinical trial imply improvement of BKPyVAN in kidney transplant recipients with an mAb-based treatment based on findings presented at the World Transplant Congress (2025) (19).

To develop safe and effective therapeutics against BKV, we generated neutralizing antibodies to BKV genotypes from VelocImmune® transgenic mice engineered to produce human antibodies (20, 21). The mice were immunized with various combinations of VP1 cDNAs to generate a broadly reactive humoral response. Hybridoma clones from the immunized mice identified monoclonal antibodies (mAbs) that broadly neutralize pseudoviruses based on BKV-I, -II, -III, and -IV genotypes. These mAbs offer therapeutic potential to limit virus spread in the host, regardless of the replicating BKV genotype present in the transplant recipient.

## METHODS AND MATERIALS

### Cells and plasmids

Human embryonic kidney (HEK)-293TT cells (ATCC # CRL-3467) were cultured in Dulbecco’s modified Eagle’s medium (DMEM, Corning, 10-013-CV) supplemented with 10% heat-inactivated fetal bovine serum (FBS), 1 mM HEPES (Corning, 25-060-CI), 100 U/mL penicillin, and 100 g/mL streptomycin (100X Pen/Strep, Corning, 30-002-CI). NIH HEK Expi293 F cells were cultured in Expi293™ Expression Medium (ThermoFisher, A1435101) and transfection systems were purchased from Invitrogen. The BKV genotypes of VP1: BKV-Ib2 (PittVR2; DQ989796), BKV-II (GBR-12; AB263920), BKV-III (KOM-3; AB211386), and BKV-IVb1 (THK-8; AB21139032); VP2 and VP2: BKV-Ib1 (KOM-5; AB211374) were cloned into mammalian expression plasmids (4, 22). MAU868 was generated from a published patent application sequence (WO 2021/252835 A1). Coding DNA sequence representative of the variable segments for the heavy and light chains were synthesized and cloned into vectors containing human G1/lambda 2 constant regions (Genscript). Heavy and light chain vectors were co-transfected in a 1:1 ratio into Expi293 cells using Expefectamine following manufacturer instructions. Supernatants were collected 5 days later, and antibody was purified using protein A HiTrap columns on an AKTA FPLC machine (Cytiva).

### BKV Pseudovirus generation

As previously described (4, 23), BKV pseudoviruses (BK-PsVs) were produced from HEK-293TT cells transfected with plasmids that express VP1, VP2, VP3, and the GFP reporter vector at ratios of 2:1:1:1 using Lipofectamine 2000 (Invitrogen). Forty-eight hours after transfection, the cells were harvested at ∼100 million cells/ml in PBS, washed 1X with PBS and lysed by addition of 0.5% Triton X-100, and the RNase A/T1 cocktail (Ambion). The lysate was incubated at 37°C overnight and cell lysates were centrifuged at 5,000×g. The BK-PsVs were isolated by ultracentrifugation through a 27–33–39% iodixanol (Optiprep, Sigma) step gradient at 300,000xg for 3.5 hrs at 16°C. The fractions from the Optiprep gradient were analyzed for detection of VP1/VP2/VP3 using SDS-PAGE. BK-PsV-transduced cells were analyzed by flow cytometry using GFP positive cells as the readout for infectivity level. Virus titer was determined based on the number of GFP-positive cells/ml of virus stock, which typically yielded ∼2.5X10^8^ infectious particles/ml.

### Mouse immunization and hybridoma fusion

VelocImmune® female mice were provided by Regeneron Pharmaceuticals and encode chimeric immunoglobulins consisting of human Vh and Vl chains and a mouse Fc domain. They can be made fully human by a simple cloning step (20, 21). The VelocImmune mice were immunized with expression plasmids for VP1, VP2 and VP3 by electroporation (see below). Briefly, mice received a prime immunization with VP1 cDNAs of BKV-I, -II, -III, or -IV genotypes with VP2 and VP3 of BKV-I genotype under different conditions (**Figure 1**), followed by a boosting immunization every 14-20 days post initial immunization. Note, we utilized genotype I VP2 and VP3 for production of all pseudoviruses based on unpublished experimental evidence that these proteins allowed for proper assembly and generation of infectious pseudovirus particles. Mice received a total of three immunizations, and blood was collected from the submandibular vein on days ∼14-20 post-priming for binding and neutralization analysis. Five mice were immunized per immunization strategy. Unexpectedly, one mouse from the Group 1 and Group 3 immunization regimens died, leaving only four animals for analysis. The animals whose sera demonstrated broad neutralization against BKV-I, -II, -III, and -IV BK-PsVs were selected for the generation of hybridomas. Final boosts with BK-PsV preparations were administered via IP injection of pseudovirus protein particles without adjuvant 3-4 days prior to harvesting the spleen. The mouse’s spleen was processed to a single-cell suspension before hybridoma generation. Splenocytes from each mouse were fused using polyethylene glycol (PEG) with hypoxanthine-aminopterin-thymidine-sensitive F23.1 cells using the ClonalCell-HY system and protocol, as described previously (StemCell Technologies) (24). Briefly, individual B-cell clones were grown on soft agar and selected for screening using a robotic ClonaCell Easy Pick instrument (Hamilton/StemCell Technologies). Individual clones were expanded, and the supernatant was used to screen for VP1 binding followed by BK-PsV neutralization.

**Figure 1.**
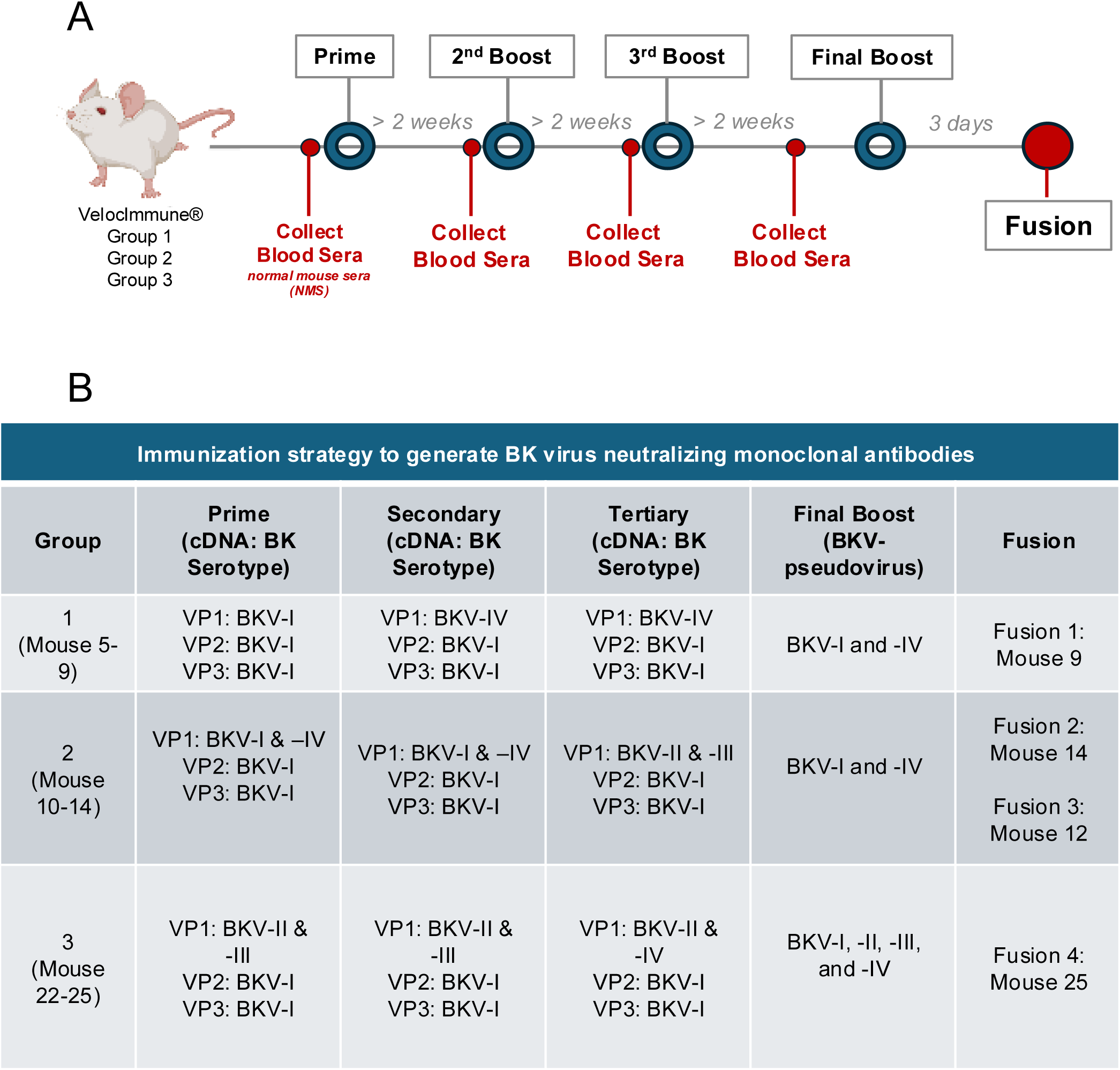
Immunization regimen for generating anti-BKV antibodies. VelocImmune® mice were immunized over a course of several months (**A**) with expression plasmids encoding BKV VP1 from genotypes I, II, III, and IV with VP2 and VP3 of BKV-I genotype in diverse combinations with a final boost of the respective BKV pseudoviruses. The specific immunization regimen was designated as Groups 1, 2, and 3 (**B**). Based on the BKV humoral response, hybridomas were generated from the spleen of designated animals (Fusions 1-4).

### Electroporation

For DNA immunization, mice were anesthetized using 4 parts ketamine HCl (100 mg/ml stock solution) to 1-part xylazine (20 mg/ml stock). Each mouse received 100 µg of plasmid DNA encoding the CDS of BK VP1, 2, or 3 reconstituted in sterile 1x PBS in a total volume of 100ul. DNA was injected subcutaneously (∼90ul) to form a skin bleb of approximately 8.7mm2. Immediately following bleb formation, a small animal, 6×6 needle electrode with 4mm spacing and 2mm depth (subQ depth) is positioned over the bleb, which is centered between the electrode rows, and pushed gently into the skin. Using the AgilePulse In Vivo Electroporation System (Harvard BTX, Harvard Biosciences, Inc.) under manufacturer-supplied pulse dynamics (Pulse Amplitude (110 V), Pulse Width (10ms), Pulse Interval (300 ms), Group Interval (500 ms), Pulse Number (8)) was used to deliver DNA to the animals following probe placement. Mice were gently placed backed into the cage and allowed to recover. Blood was collected 2 weeks after the last immunization and used for analysis. The pre-immunization bleed sera (pre-bleed or normal mouse sera, NMS) served as the negative control.

### Hybridoma supernatant immunoglobulin binding assay

Expi293F cells were transfected with the expression plasmids encoding VP1 protein of the BKV-I genotype. Forty-eight hours post-transfection, cells were permeabilized with BD Cytofix/Cytoperm (BD #554722), washed, and stained with hybridoma clone supernatant (1:2) in PBS with 0.5% BSA for 1 hr. After 2 washes in PBS/BSA, cells were incubated with FITC-conjugated goat anti-mouse IgG secondary (Jackson Immunoresearch) for 30 mins. Cells were washed once in PBS and resuspended in 50 µl PBS to be analyzed on a high throughput flow cytometry (HTFC) device (Intellicyt). The mean fluorescence intensity (MFI) of each sample was measured based on the analysis of 10,000 events/sample. The MFI was normalized to Relative MFI based on GFP signal from each well over background MFI using conditioned media. The Relative MFI was plotted on using a heat map for easy comparison among clones.

### BK-PsV infectivity assay

Infectivity assay of BKV-I, -II, -III, and -IV BK-PsV was performed as follows: 15,000 HEK-293TT cells were plated in each well of 96 well plate. The next day, the hybridoma supernatant or purified antibodies were diluted in DMEM, respectively, and mixed with BKV-I, -II, -III, and -IV BK-PsV to yield ∼30% infection (multiplicity of infection (moi) 0.3). The antibody/BK-PsVs mixtures were added to each well of cells and infection was evaluated by flow cytometry using GFP fluorescence as the readout using the HTFC (Intellicyt) 24 hours post-infection (hpi). The population of infected cells was validated by an increase in mean fluorescent intensity (MFI) compared to non-infected cells. The % GFP (+) cells were determined for each sample, and the relative percentage (%) infection was calculated based on untreated/control-treated BK-PsV infection as 100% infection.

### Antibody sequence analysis of heavy chain variable region and junctional diversity

Sequencing of variable heavy and kappa chains was obtained using SMARTer 5’ RACE technology (Takara Bio USA) adapted for immunoglobulins to amplify the variable genes from the heavy and kappa chains. Briefly, RNA was extracted from each hybridoma using a RNeasy Mini Kit (Qiagen, 74004), followed by first-strand cDNA synthesis using constant gene-specific 3’ primers (GSP1) based on the specific mouse isotype of the hybridoma and incubation with the SMARTer II A Oligonucleotide and SMARTscribe reverse transcriptase (Takara, 634858). [GSP1 Primers (5’-3’): mG1-AGAGGTCAGACTGCAGGACA, mG2a-CTTGTCCACTTTGGTGCTGC, mG2b-GACAGTCACTGAGCTGCTCA, mG2b-GACAGTCACTGAGCTGCTCA, mcK-CCAACTGTTCAGGACGCCAT]. PCR amplification of the first-strand cDNA product was then performed using SeqAmp DNA Polymerase (Takara, 638504) with a nested 3’ primer (GSP2 Primer) to the constant genes and a 5’ universal primer based on universal primer sites added to the 5’ end during cDNA generation. [GSP2 Primers (5’-3’): mG1-CCCAGGGTCACCATGGAGTT, mG2a-GGTCACTGGCTCAGGGAAAT, mG2b-CTTGACCAGGCATCCCAGAG, mG3-GACAGGGCTCCATAGTTCCATT, mCk-CTGAGGCACCTCCAGATGTTAAC] Purified PCR products were submitted for Sanger sequencing using 3’ constant gene primers (GeneWiz, South Plainfield, NJ). Using BLAST, sequence results were analyzed against the IMGT human databank of germline genes using V-Quest (http://imgt.org/).

### Isotyping and monoclonal antibody purification

Isotyping for the constant gene of the antibodies was performed with the Mouse Immunoglobulin Isotyping Kit (BD, #550026) as per the manufacturer’s protocol. Monoclonal antibodies were purified by FPLC on an ÄKTA pure FPLC system on protein G affinity columns (HiTrap-1mL, GE/Cytiva, #17-0404-01) followed by dialysis against PBS and quantification by BCA and OD at 280 nm. The purified monoclonal antibodies reagents will be made available with the scientific community upon execution of a Material Transfer Agreement between the respective institutions.

### Quantitation of monoclonal antibodies

Immunoglobulins in the hybridoma supernatant were quantitated using label-free, biolayer interferometry (BLI) on an Octet Red96 (ForteBio, Sartorius). Briefly, optically capable biosensors conjugated with protein G were incubated with hybridoma supernatants containing mAbs. Supernatants were measured undiluted and diluted 1:10 in conditioned media and compared to an isotype standard, diluted in conditioned media in a 1:2 dilution series ranging from 1.56 μg/ml to 100 μg/ml. Standard curves were analyzed in ForteBio Data Analysis Software v.10 (ForteBio) using a 5-parameter logistics (5PL) dose response curve fitting model on the initial binding slopes. MAb concentrations were then calculated from the standard curve. Diluted samples were compared to undiluted samples after the application of the respective dilution factors. Total concentrations were averaged together from the diluted and undiluted samples.

### Statistics

The statistical tests were performed using GraphPad Prism 9 software (La Jolla, CA). The half (50%)-maximal inhibitory concentration (IC50) and IC80 values were calculated using three-parameter non-linear regression analysis (GraphPad), after antibody concentrations (x-axis) were transformed to log scale.

### Ethical approval

Animals were housed in an AAALAC (Association for the Assessment and Accreditation of Laboratory Animal Care) approved facility. The animals were provided with enrichment and cared for by qualified animal technicians. All animal studies were approved by the Icahn School of Medicine at Mount Sinai’s Institutional Animal Care and Use Committee (IACUC), IACUC # IPROTO202200000097, and the animal studies were strictly adhered to ARRIVE guidelines.

## RESULTS

### Humoral response of VelocImmune® mice immunized with BKV genotypes

Despite the high degree of sequence similarity of VP1 among the four genotypes (>95% (18)), a humoral response against one genotype does not necessarily protect against a different genotype (17). We sought to generate broadly neutralizing human monoclonal antibodies against the viral surface protein VP1 of BKV genotypes for future therapeutics against BKV-associated diseases. To that end, VelocImmune^®^ mice (21) were immunized with diverse combinations of cDNAs encoding the VP1 proteins of the BKV-Ib2, -II, -III, and -IVb1 genotypes that included BKV-Ib1 VP2 and VP3 to generate an infectious virus particle due to a native VP1 molecule (**Figure 1A**). The subtypes of the BKV-I and BKV-IV genotypes will be abbreviated BKV-I and BKV-IV for simplicity. Using cDNAs as the primary immunogen allows for prolonged expression of the entire protein to enhance immune recognition, including post-translational modifications (25, 26). Three immunization regimens (Group 1, 2, and 3) were utilized to promote a broadly reactive humoral response to all BKV VP1 genotypes (**Figure 1B**). The sera from the respective immunized animals were collected to evaluate the humoral response to VP1 of BKV-I, -II, -III, and -IV (**Figure 2**). The diluted sera (1:200, 1:1,000, and 1:5,000) were analyzed for neutralization activity with the infectivity assay against BKV-I, -II, -III, and -IV BK-PsV expresses GFP, a proxy for a BKV particle infectivity. The BK-PsV transduced HEK-293TT cells were analyzed for GFP positivity 24 hours post-infection (hpi) by flow cytometry. The percentage of GFP positive cells was determined and normalized to relative percent infection using BK-PsV incubated with normal mouse serum as 100% relative infection.

**Figure 2.**
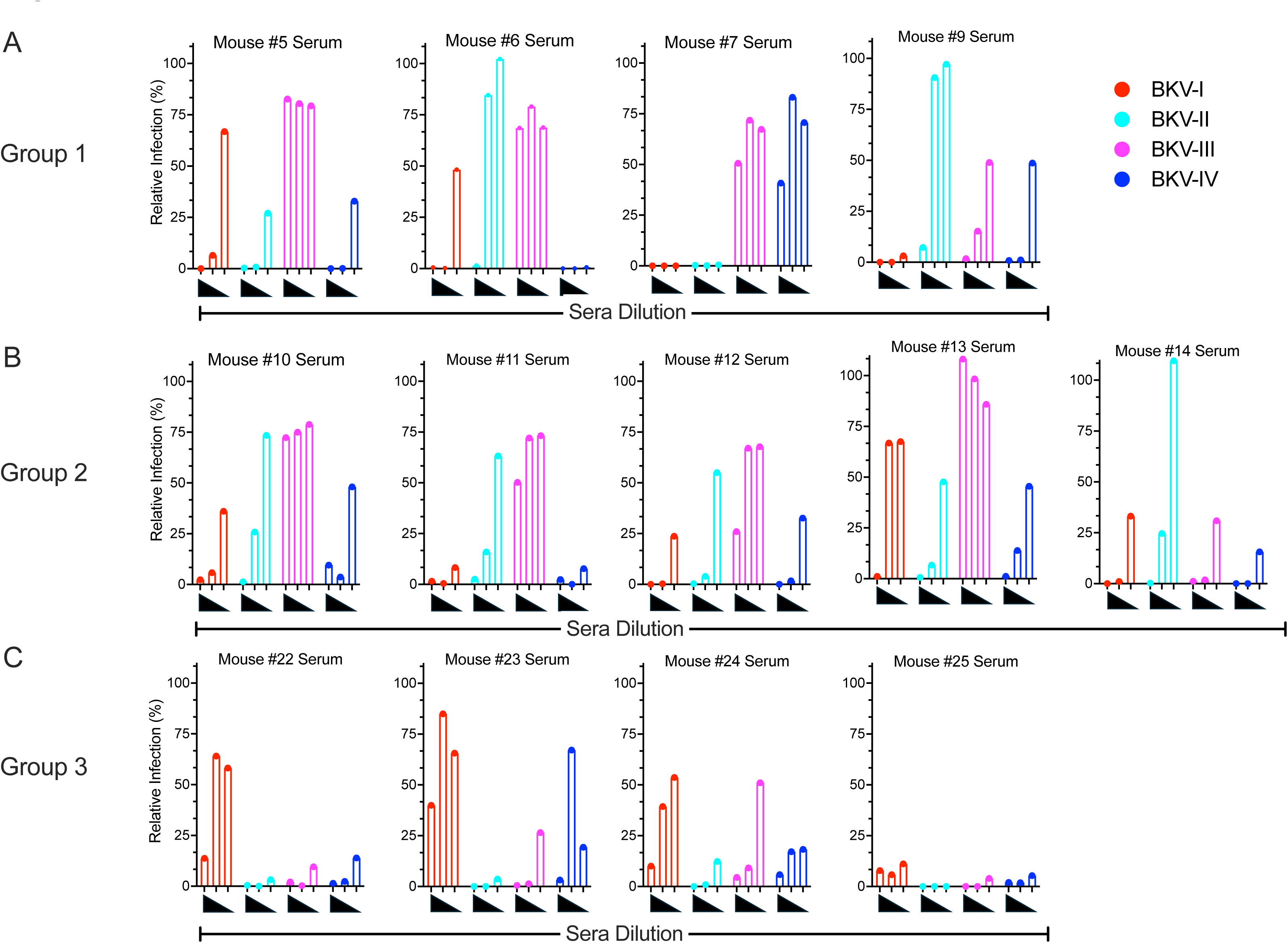
Neutralization activity of sera from VelocImmune® mice immunized with BKV antigens. The sera from VelocImmune® mice immunized with BKV antigen combinations (Group 1 (**A**), Group 2 (**B**), Group 3 (**C**)) were evaluated for neutralization of BK-PsV-I, -II, -III, and -IV expressing GFP. The diluted sera (1:200, 1:1,000, and 1:5,000) from the designated mouse was pre-incubated with BK-PsVs and added to HEK-293TT cells. BK-PsV infection was evaluated 24 hours post infection (hpi) using flow cytometry to determine the % of infected cells based on GFP positivity. The relative infection (%) was determined using BK-PsV treated with normal mouse sera as 100%.

The neutralization serology profiles of the immunized animals were distinct among the immunization regimens. In all cases, the sera from the immunized mice inhibited at least two BKV genotypes in a dilution-dependent manner. The serum from Group 1 immunized animals with BKV-I and -IV VP1 (Mouse (Ms)-5, -6, -7, and -9) were effective at inhibiting BK-PsV-I and BK-PsV-IV infectivity (**Figure 2A**). Sera from Ms-5, Ms-6, and Ms-7 were unable to block BK-PsV-III infection, while Ms-9 sera demonstrated inhibition of all BK-PsV genotypes. For the Group 2 immunized animals (Ms-10, -11, -12, -13, -14), only Ms-12 and Ms-14 sera showed concentration-dependent neutralization of all BK-PsV genotypes (**Figure 2B**). Strikingly, the Group 3 animals (Ms-22, -23, 24, -25), immunized with VP1 cDNA of BKV-II and BKV-III followed by BK-PsV of all genotypes, demonstrated a broad and robust neutralization response against all four BKV genotypes (**Figure 2C**). We suspect that the humoral responses against BKV-II and BKV-III VP1 target conserved domains within the VP1 protein that are also present in the BKV-I and BKV-IV VP1 molecules. The result is reminiscent of prior work by Pastrana and colleagues suggesting that some BKV genotypes (particularly BKV-II) may be better at eliciting cross-neutralizing antibody responses (4). Collectively, these studies show the immunogenicity of BKV VP1 protein can inhibit infection of diverse BKV genotypes.

### Generation and identification of anti-BKV monoclonal antibodies

Mice Ms-9, Ms-12, Ms-14, and Ms-25, whose sera demonstrated cross-neutralization activity against BKV-I, -II, -III, and -IV BK-PsV genotypes, were selected for the generation of hybridomas to identify broadly reactive immunoglobulin clones. The supernatants from the hybridoma clones were screened for binding to BKV-I VP1 using a flow cytometry binding assay of HEK-293 cells transfected with BKV-I VP1 cDNA (**Figure 3A-D**). The hybridoma clone supernatants (1:2 dilution) were incubated with BKV-I VP1-transfected HEK-293 cells followed by anti-mouse IgG-FITC and then analyzed for mean fluorescence intensity (MFI) on a flow cytometer. Controls included normal mouse sera from non-immunized mice and polyclonal sera from the immunized animals. The immunization strategy included VP2 and VP3 cDNA and protein and thus some of the clones may be specific for VP2 and VP3.

**Figure 3.**
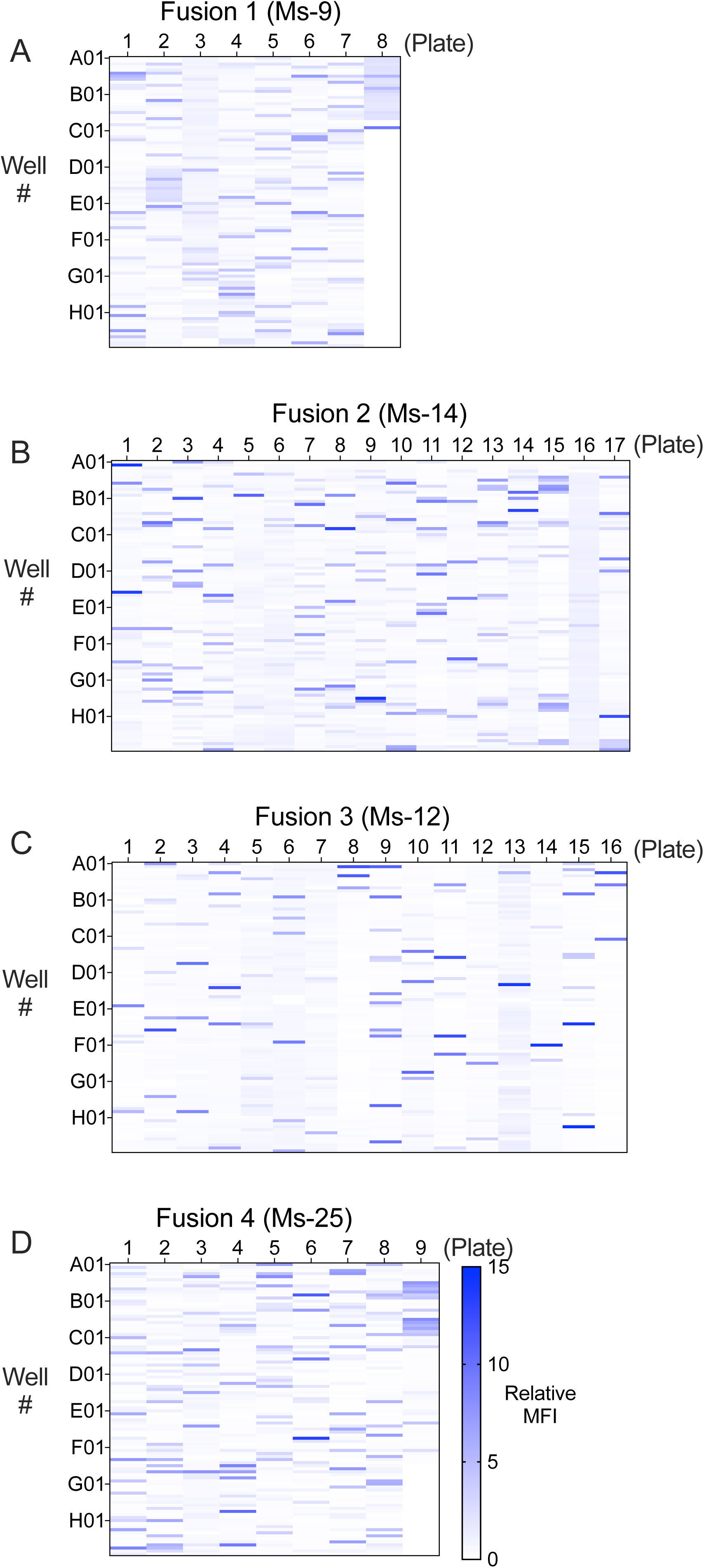
Binding profile of hybridoma clones from BKV-immunized VelocImmune® mice. Hybridoma clones generated from BKV-immunized VelocImmune® mice were analyzed for binding to VP1 BKV-I transfected HEK-293 cells at 48 hrs post-transfection. The hybridoma supernatants (1:2) from Fusion 1 (Ms-9) (**A**), Fusion 2 (Ms-14) (**B**), Fusion 3 (Ms-12) (**C**), and Fusion 4 (Ms-25) (**D**) were incubated with permeabilized transfected cells and evaluated by flow cytometer. The mean fluorescence intensity (MFI) was measured from each sample and normalized to fold change over background MFI to define a Relative MFI (0–15). The Relative MFI was plotted as a heatmap for each clone in which the X-axis is the 96-well plate number and the Y-axis indicates the 96 well plate number (A1-H12).

However, we selected to focus on the clones that can neutralize BKV through targeting of the virus surface protein VP1. Hybridoma clones from Ms-9 (Fusion 1, 693 clones), Ms-12 (Fusion 2, 1628 clones), Ms-14 (Fusion 3, 1443 clones), and Ms-25 (Fusion 4, 823 clones) were evaluated for binding to VP1-transfected cells in which the MFI of each clone was normalized as fold change over background and represented as Relative MFI. Clones that demonstrated a 2-fold increase in relative MFI (Fusion 1: Ms-9, 107 clones; Fusion 2: Ms-14, 173 clones; Fusion 3: Ms-12, 154 clones; Fusion 4: Ms-25,109 clones) were selected for isotype analysis. The anti-BKV VP1 binding clones were subjected to isotype analysis to identify IgG1, IgG2a, and IgG2b immunoglobulins. IgM and IgG3 isotypes were excluded because they are more prone to proteolysis and aggregation in solution, respectively, and are generally considered less viable for development as therapeutics (27). Based on IgG isotype and hybridoma viability, 353 total binding clones (Fusion 1: Ms-9, 75 clones; Fusion 2: Ms-14, 96 clones; Fusion 3: Ms-12, 123 clones; Fusion 4: Ms-25, 59 clones) were selected for analysis in the BK-PsV infectivity assays.

### Analysis of neutralizing anti-BK virus mAbs

The VP1 binding clones were next evaluated for neutralization of BKV-I, -II, -III, or -IV via the pseudovirus infectivity assay (**Figure 4**). Neutralizing clones to BK-PsV-I were highly representative in the Group 1 and 2 immunization regimens (Fusion 1, (Ms-9); Fusion 2 (Ms-14); Fusion 3 (Ms-12)), with 8 clones reducing infectivity of all four BK-PsV genotypes (**Figures 4 and 5A-C**). Strikingly, clones from the Group 3 immunization regimen (Fusion 4, Ms-25) were quite effective at limiting infectivity of BK-PsV-II and -III, with many clones reducing infectivity of all genotypes. The Group 3 immunization strategy generated the highest number (28 clones) of broadly neutralizing BK-PsV clones (**Figures 4 and 5D**), a result consistent with the binding profile of the clones (**Figure 3**). Overall, 36 clones (Figure 4, highlighted in bold) effectively neutralized (>75%) BK-PsV-I, -II, -III, and -IV while 184 clones blocked a single BK-PsV genotype, mostly BKV-I (**Figure 5A-D**). The specificity for BKV-I may be due to the expression levels of BKV-I VP1 cDNA or immunodominant domains within BKV-I VP1. The data suggest that varying immunogens and order of immunization may play a significant role in creating a broadly reactive humoral response.

**Figure 4.**
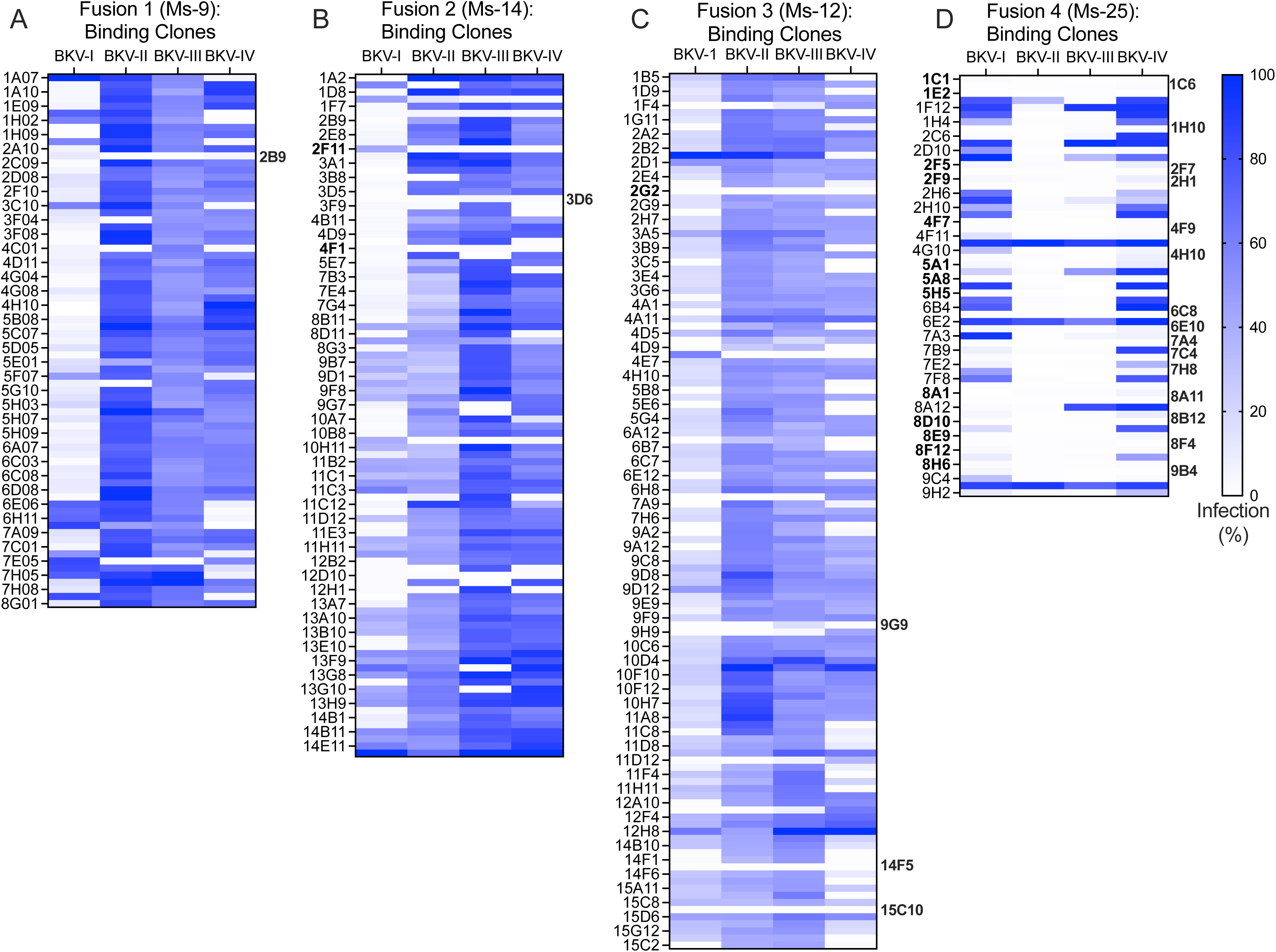
Neutralization capacity of VP1-binding clones to limit infection of BK-PsV genotypes. The BKV-I VP1 binding clones were evaluated in BK-PsV-infectivity assays. The hybridoma supernatant of the VP1-binding clones from Fusion 1 (Ms-9, **A**), Fusion 2 (Ms-14, **B**), Fusion 3 (Ms-12, **C**), and Fusion 4 (Ms-25, **D**) was pre-incubated (1:500) with the indicated BK-PsV genotype, added to HEK-293TT cells, and evaluated 24hpi using flow cytometry to determine the % of GFP positive cells. The relative infection % was determined using BK-PsV incubated with media as 100%. The relative % infection was plotted for each clone (Y axis) as a heat map for each BK-PsV genotype. The clones in bold are the broadly neutralizing monoclonal antibodies.

**Figure 5.**
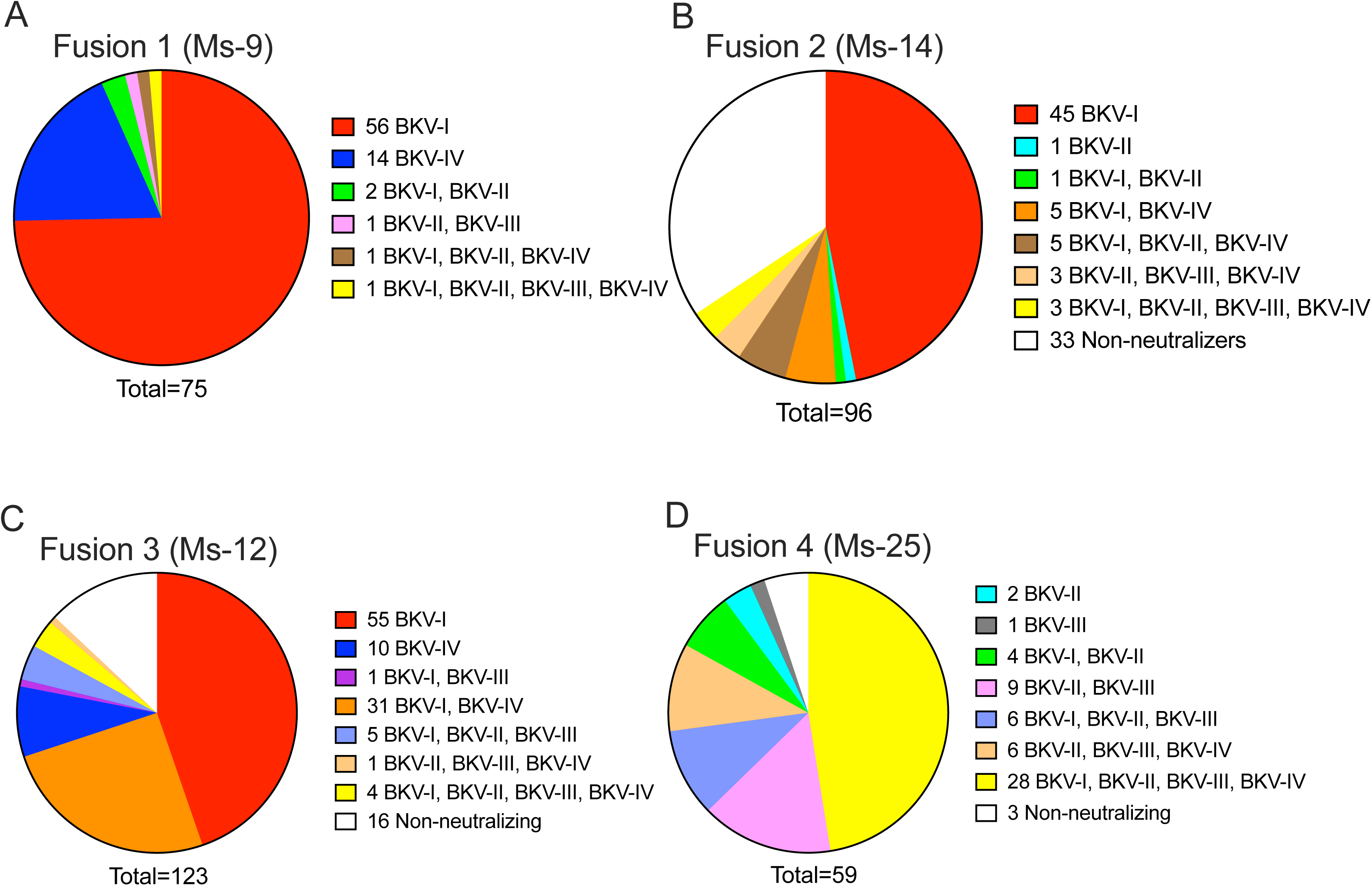
Summary of neutralization results of VP1 binding hybridoma clones. Based on the neutralization data in Figure 4, the VP1-binding hybridoma clones with <75% infectivity from Fusion 1 (Ms-9, **A**), Fusion 2 (Ms-14, **B**), Fusion 3 (Ms-12, **C**), and Fusion 4 (Ms-25, **D**) were categorized into selective inhibition of BK-PsV genotypes.

Based on their neutralization profile and ability to limit infection in all four BK-PsV genotypes, 8 anti-BKV clones from Fusions 1, 2, and 3, along with 15 of the 28 clones from Fusion 4, were selected for nucleotide sequencing of their immunoglobulin variable segment (**Table S1**). Sequence analysis was utilized to eliminate identical clones and categorize the remaining unique clones into clonotype families. Clonotype families were defined by heavy sequences that shared homologous CDR3 regions (exact length and >95% nucleotide identity) with shared V(D)J gene usage and homologous light chain pairs (light chain CDR3 homology and VJ gene usage). The clones averaged 5-8% nucleotide mutations in their V-gene segments when compared to the closest germline gene by BLASTing the sequences against the IMGT human germline databank, indicative of affinity-matured, somatically hypermutated antibodies. Based on their sequences and mutation frequencies, we proposed that these antibodies evolved through affinity maturation by our immunization strategy to improve their specificity and binding affinity (28, 29). Collectively, we identified six diverse clonotype families (A-F) based on CDR3 and heavy gene sequences (**Table S1**) representing antibodies that broadly limit infection of BK-PsV-I, -II, - III, and -IV.

### Evaluation of anti-BKV mAb neutralization

We next evaluated the neutralization capacity of the genetically distinct and cross-reactive anti-BKV mAbs (**Table S1**) using a BK-PsV infectivity assay. Using hybridoma supernatant, the clones’ concentrations were determined and compared to isotype-matched standard controls with label-free Bio-Layer Interferometry (BLI) on an Octet Red 96. The supernatants from twelve clones representing all six clonotypes were examined for their ability to neutralize infection of the BK-PsV-I, -II, -III, and -IV genotypes (**Table 1**). The approximate inhibitory concentrations at 50% (IC50) of the mAb clones across all four BK-PsV genotypes were determined based on % of GFP positive cells. The clones exhibited a wide range of IC50 values: 9-667 pM (BK-PsV-I), 8->4000 pM (BK-PsV-II), 8-350 pM (BK-PsV-III), and 10-148 pM (BK-PsV-IV). Notably, many clones --2B9, 3D6, 4F1, and 9G9--did not meet the threshold of IC50 <100 pM for one or more genotypes (**Table 1**) and hence were excluded from further analysis. These results demonstrate the identification of several genetically unique mAbs that broadly neutralize BKV-I, -II, -III, and -IV genotypes.

**Table 1.**
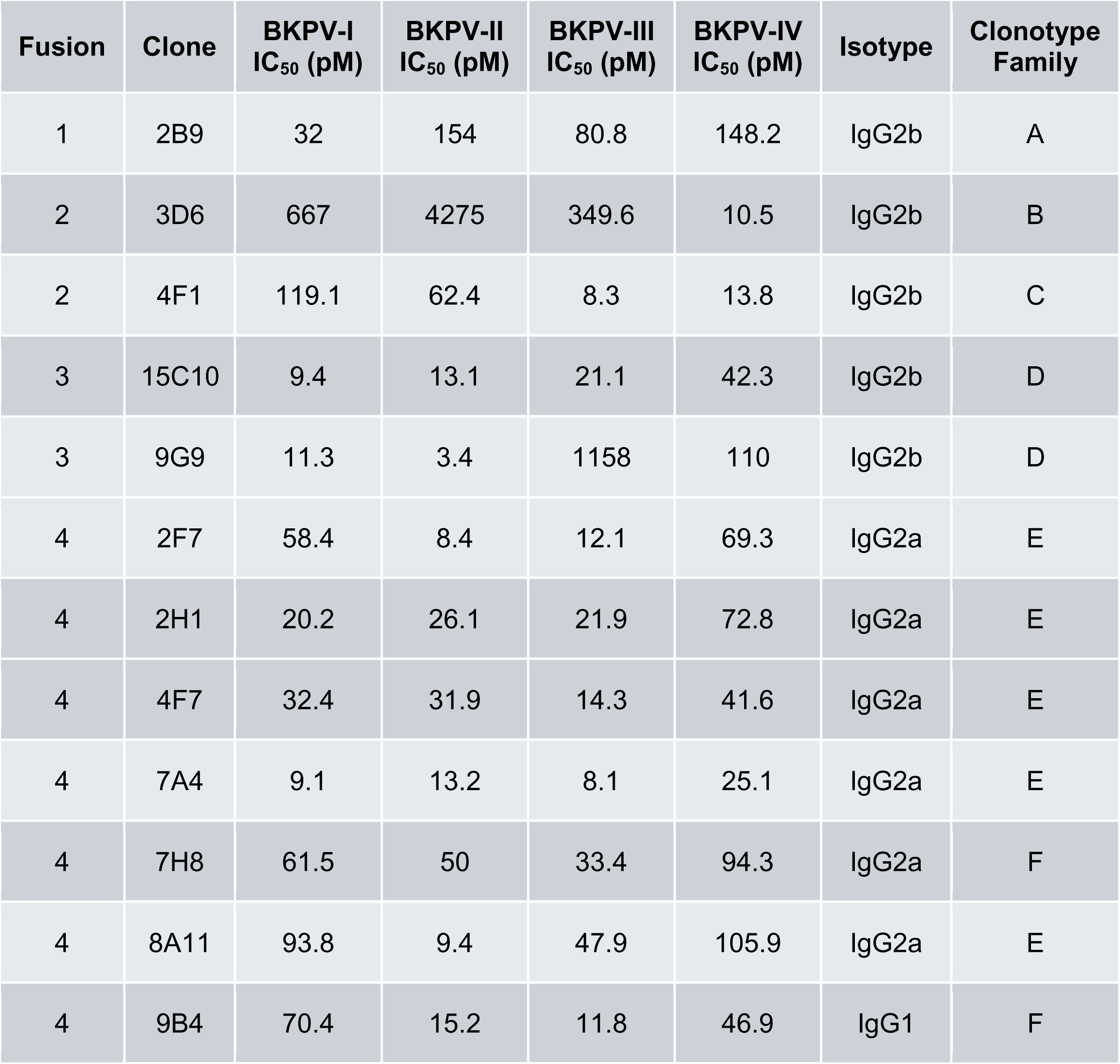
Summary of IC_50_ (pM) values of anti-BKV neutralizing monoclonal antibodies

| **Fusion** | **Clone** | **BKPV-I**<br>**IC<sub>50</sub> (pM)** | **BKPV-II**<br>**IC<sub>50</sub> (pM)** | **BKPV-III**<br>**IC<sub>50</sub> (pM)** | **BKPV-IV**<br>**IC<sub>50</sub> (pM)** | **Isotype** | **Clonotype**<br>**Family** |
| --- | --- | --- | --- | --- | --- | --- | --- |
| 1 | 2B9 | 32 | 154 | 80.8 | 148.2 | IgG2b | A |
| 2 | 3D6 | 667 | 4275 | 349.6 | 10.5 | IgG2b | B |
| 2 | 4F1 | 119.1 | 62.4 | 8.3 | 13.8 | IgG2b | C |
| 3 | 15C10 | 9.4 | 13.1 | 21.1 | 42.3 | IgG2b | D |
| 3 | 9G9 | 11.3 | 3.4 | 1158 | 110 | IgG2b | D |
| 4 | 2F7 | 58.4 | 8.4 | 12.1 | 69.3 | IgG2a | E |
| 4 | 2H1 | 20.2 | 26.1 | 21.9 | 72.8 | IgG2a | E |
| 4 | 4F7 | 32.4 | 31.9 | 14.3 | 41.6 | IgG2a | E |
| 4 | 7A4 | 9.1 | 13.2 | 8.1 | 25.1 | IgG2a | E |
| 4 | 7H8 | 61.5 | 50 | 33.4 | 94.3 | IgG2a | F |
| 4 | 8A11 | 93.8 | 9.4 | 47.9 | 105.9 | IgG2a | E |
| 4 | 9B4 | 70.4 | 15.2 | 11.8 | 46.9 | IgG1 | F |

To further characterize the BKV neutralizing mAbs, select clones across different clonotype families were chosen and cloned with a human IgG1 Fc domain and evaluated for neutralization of BK-PsVs. Clones 2H1 (Family E), 7A4 (Family E), 7H8 (Family F), and 15C10 (Family D) were selected due to their IC50 values in the BK-PsV infectivity assays (**Table 1**). In addition, a previously identified anti-BKV mAb that targets VP1, MAU868 (30), was examined to compare the neutralization profiles of the newly characterized anti-BKV mAbs (**Figure 6A-D**). Clones 2H1, 7A4, 7H8, and 15C10 demonstrated robust neutralization of all BK-PsV genotypes with IC50 values of 7-50 pM for BK-PsV-I, 12-80 pM for BK-PsV-II, 45-112 pM for BK-PsV-III, and 85-243 pM for BK-PsV-IV and IC80 values 13-104 pM for BK-PsV-I, 15-140 pM BK-PsV-II, 84-330 pM for BK-PsV-III, and 148-280 pM for BK-PsV-IV (**Table 2**).

**Figure 6.**
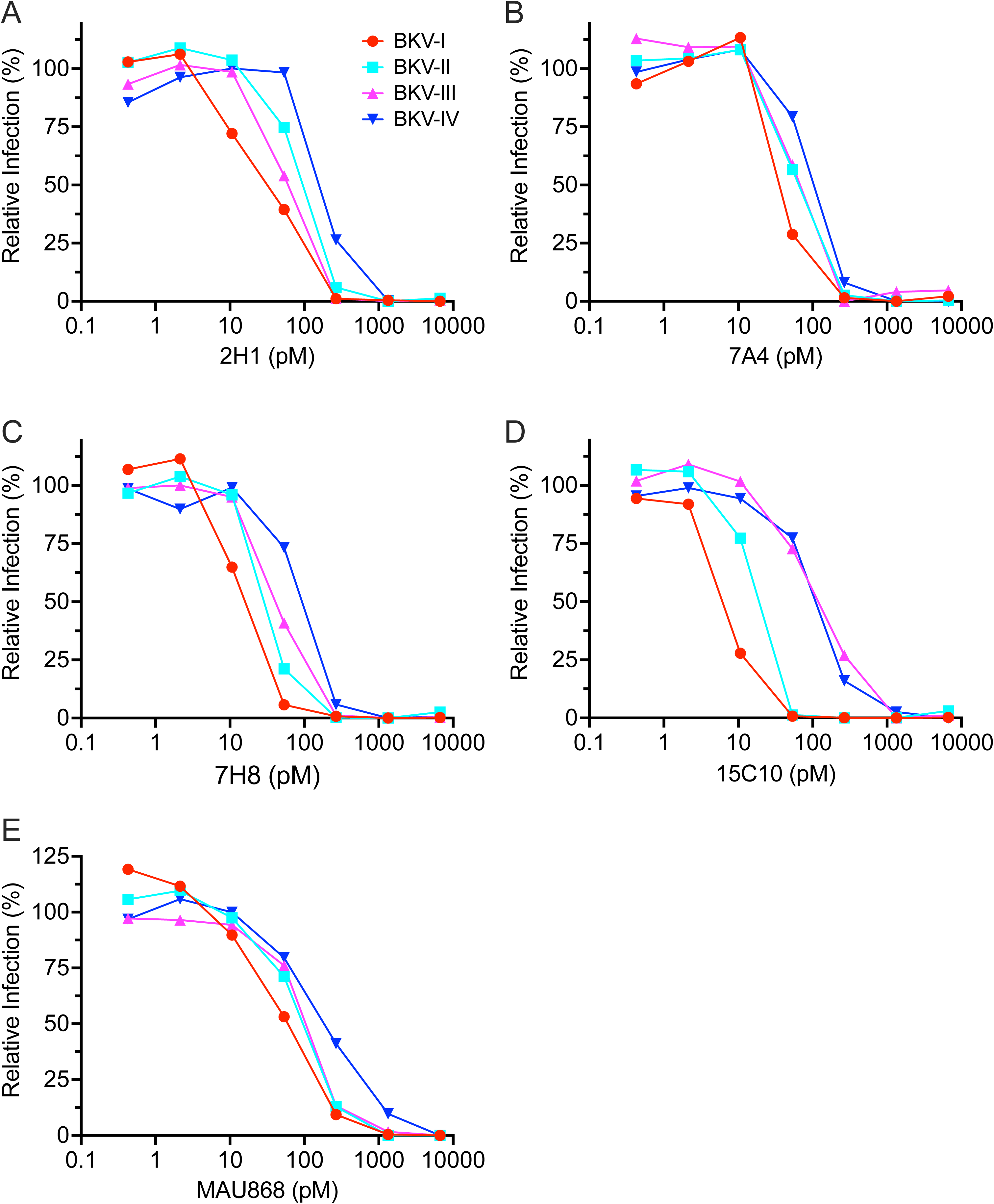
Analysis of broadly neutralizing human mAbs against BKV genotypes. Analysis of human anti-BKV mAbs in the BK-PsV-I, -II, -III, and -IV infectivity assays was performed with the anti-BKV mAb clones 2H1 (**A**), 7A4 (**B**), 7H8 (**C**), 15C10 (**D**), and MAU868 (**E**). MAbs (0-6670 pM) pre-incubated with BK-PsV were used to infect HEK-293TT cells and evaluated 24 hpi for % of GFP positive cells. The relative infection (%) was determined using BK-PsV pre-incubated with non-BKV mAb as 100%.

**Table 2:** Summary of IC_50_ and IC_80_ values (pM) of fully human anti-BKV mAbs

|  | BKV-I |  | BKV-II |  | BKV-III |  | BKV-IV |  |
| --- | --- | --- | --- | --- | --- | --- | --- | --- |
| Clone | IC<sub>50</sub><br>(pM) | IC<sub>80</sub><br>(pM) | IC<sub>50</sub><br>(pM) | IC<sub>80</sub><br>(pM) | IC<sub>50</sub><br>(pM) | IC<sub>80</sub><br>(pM) | IC<sub>50</sub><br>(pM) | IC<sub>80</sub><br>(pM) |
| 2H1 | 30 | 104 | 78.5 | 139 | 56.3 | 84.2 | 243 | 280 |
| 7A4 | 48.3 | 55.5 | 54.1 | 63.9 | 53.7 | 63.5 | 89.1 | 160.8 |
| 7H8 | 12.5 | 22.5 | 32.8 | 53.8 | 45.4 | 85.1 | 85.2 | 148 |
| 15C10 | 7.2 | 13.1 | 12.5 | 15.2 | 112 | 331 | 109 | 231 |
| MAU868 | 35 | 170 | 82.3 | 203 | 104 | 211 | 189 | 686 |

Monoclonal antibody 7A4 and 7H8 were the most effective neutralizers across all BKV genotypes (**Figure 6B and C**, **Table 2**). In fact, the Hill slope coefficient is steeper for clones 7A4 (2-10 fold increase of the Hill slope) and 7H8 (2-fold increase of the Hill slope) compared to MAU868. For antiviral agents, steeper Hill slopes are considered more favorable for suppressing the emergence of resistance mutations (31).

Interestingly, compared to the purified antibody, some clones evaluated from the hybridoma supernatant showed lower IC50 values; perhaps due to cloning of the human IgG1 Fc domain or mAb concentration being underestimated (32, 33). Notably, the IC50 values of the newly characterized neutralization mAbs were comparative or lower than MAU868 supporting their effectiveness as a potential VP1 target treatment. Collectively, these studies demonstrate the proof of concept that mAbs targeting VP1 of the BK virus can broadly limit infection of all genotypes.

## DISCUSSION

We have identified a panel of genetically distinct broadly reactive mAbs that target the VP1 of BKV-I, -II, -III, and -IV genotypes and effectively block the infectivity of cognate pseudoviruses (BK-PsV). Specifically, clones from four genetically diverse mAb genotype families effectively reduced infection with BKV-I, -II, -III, and -IV BK-PsV (**Figure 6**). Importantly, clones 7A4 and 7H8 demonstrated relatively low IC50 values in comparison to a previously characterized BKV neutralizing mAb, MAU868, undergoing clinical trials (Clinical trial#NCT04294472) (**Table 2**). Further, we identified numerous neutralizing clones specific for a single BKV genotype for future utilization as genotype-specific biologic tools and/or therapeutics in possible combination studies. Collectively, we have identified new human mAbs that interact with the VP1 protein of different BKV genotypes and neutralize their associated pseudoviruses. These mAbs represent potential therapeutic candidates for limiting BKV-associated diseases by preventing virus infection and spreading in at-risk kidney transplant recipients. Consistent with our findings, an anti-BKV VP1 mAb has been recently characterized to limit infection of all BKV genotypes (34).

Neutralizing mAbs were screened based on binding to different VP1 BKV genotypes and inhibiting transduction of different BK-PsVs genotypes. The immunization regimen generated neutralizing mAbs against VP1, despite the high sequence identity (>95%) among the different genotypes. The VP1 amino acid sequence identity among the genotypes is ∼95% except for the BC loop (amino acids 57-89), which has only ∼65% homology. Cross-reactive neutralizing clones likely target common epitopes within the four genotypes to limit virus infection. The cross-reactive mAb MAU868 was demonstrated to interact with VP1 through the contact residues of Y169, R170, and K172 (30). Structural predications of the cross-reactive clones 2H1, 7A4, 7H8, and 15C10 generated with ColabFold modeling (Supplemental Figure 1) illustrates a structurally distinct binding pocket of the Fab domain than MAU858, suggesting that these newly identified BKB mAbs bind a epitope distinct from MAU868. -In the future, it will be useful to see whether the hypothetical common epitopes are conserved in BKV’s close relative, JC polyomavirus.

Mouse 25 of Group 3, initially immunized with BKV-II and -III VP1 followed by BKV-I and -IV VP1, generated the largest percentage of cross-reactive mAbs suggesting a possible role of immunization order in generating broadly specific anti-BKV antibodies. In contrast, primary immunizations with BKV-I and BKV-IV VP1 genotypes mostly generated mAbs to BKV-I and -IV with only a small number being cross-reactive. We can therefore infer that additional factors, beyond primary sequence, influence VP1 immunogenicity. Specifically, the overall level of VP1 expression and proper folding into its native pentameric complexes may play an important role in developing a broad humoral response to immunogenic regions of VP1. The VP1 from the BKV-II and -III appears to expose immunodominant regions of VP1, allowing for the generation of antibodies to conserved epitopes among the VP1 protein of all BKV genotypes.

Due to their polyclonal nature, intravenous immunoglobulins (IVIGs) can neutralize all four BKV genotypes *in vitro* and increasing BKV antibody titers *in vivo* when treating patients (36–38). Clinically, IVIG administration, particularly alongside immunosuppression reduction, led to a reduction of BKV viral load in 60–86% of treated patients (39–41). Numerous studies have reported that IVIG administration under different conditions of immunosuppression and in combination with other drugs have reported lower BKV-DNAemia (42–47). These studies support the overall premise that specific anti-BKV antibodies may have protective value against BKV viral load. Several Phase 3 trials (NCT05325008 and NTC04222023) analyzing IVIG treatment under conditions of BKV-DNAemia and other transplant factors, respectively (18), are being pursued. Similarly, monoclonal antibody, MAU868 (Clinical trial#NCT04294472), completed a Phase 2 trial in which subjects had a 1.14 log_10_ decrease (p=0.051) in BKV viral load but no change in the estimated glomerular filtration rate (48). In addition, the evaluation of mAb MTX-005 (Clinical trial NCT0569582) in a Phase 2/3 trial examined BKV-DNAemia in kidney transplant patients (18). . These studies provide potential intervention strategies to increase the neutralizing antibody titer to reduce BKV-DNAemia, thereby potentially decreasing the incidence of BKVAN. Thus, our newly identified broadly neutralizing mAbs can be developed into new therapeutics to limit BKV-associated diseases.

## Supporting information

Text is within the figure

## Acknowledgements

We thank Regeneron Pharmaceuticals and NIH Institutional Research Awards - AI R56174062, AI R01139258, and AI R56174062 for supporting these studies.

## REFERENCES

1. Zhou X, Zhu C, Li H. 2023. BK polyomavirus: latency, reactivation, diseases and tumorigenesis. Front Cell Infect Microbiol 13:1263983.

2. Helle F, Brochot E, Handala L, Martin E, Castelain S, Francois C, Duverlie G. 2017. Biology of the BKPyV: An Update. Viruses 9:327.

3. Stroh LJ, Stehle T. 2014. Glycan Engagement by Viruses: Receptor Switches and Specificity. Annu Rev Virol 1:285–306.

4. Pastrana DV, Ray U, Magaldi TG, Schowalter RM, Cuburu N, Buck CB. 2013. BK polyomavirus genotypes represent distinct serotypes with distinct entry tropism. J Virol 87:10105–13.

5. Geoghegan EM, Pastrana DV, Schowalter RM, Ray U, Gao W, Ho M, Pauly GT, Sigano DM, Kaynor C, Cahir-McFarland E, Combaluzier B, Grimm J, Buck CB. 2017. Infectious Entry and Neutralization of Pathogenic JC Polyomaviruses. Cell Rep 21:1169–1179.

6. Zhong S, Randhawa PS, Ikegaya H, Chen Q, Zheng HY, Suzuki M, Takeuchi T, Shibuya A, Kitamura T, Yogo Y. 2009. Distribution patterns of BK polyomavirus (BKV) subtypes and subgroups in American, European and Asian populations suggest co-migration of BKV and the human race. J Gen Virol 90:144–52.

7. Dalianis T, Hirsch HH. 2013. Human polyomaviruses in disease and cancer. Virology 437:63–72.

8. Leuzinger K, Naegele K, Schaub S, Hirsch HH. 2019. Quantification of plasma BK polyomavirus loads is affected by sequence variability, amplicon length, and non-encapsidated viral DNA genome fragments. J Clin Virol 121:104210.

9. Myint TM, Chong CHY, Wyld M, Nankivell B, Kable K, Wong G. 2022. Polyoma BK Virus in Kidney Transplant Recipients: Screening, Monitoring, and Management. Transplantation 106:e76–e89.

10. Nourie N, Boueri C, Tran Minh H, Divard G, Lefaucheur C, Salmona M, Gressens SB, Louis K. 2024. BK Polyomavirus Infection in Kidney Transplantation: A Comprehensive Review of Current Challenges and Future Directions. Int J Mol Sci 25.

11. Kotton CN, Kamar N, Wojciechowski D, Eder M, Hopfer H, Randhawa P, Sester M, Comoli P, Tedesco Silva H, Knoll G, Brennan DC, Trofe-Clark J, Pape L, Axelrod D, Kiberd B, Wong G, Hirsch HH, Transplantation Society International BKPCG. 2024. The Second International Consensus Guidelines on the Management of BK Polyomavirus in Kidney Transplantation. Transplantation 108:1834–1866.

12. Starrett GJ, Yu K, Golubeva Y, Lenz P, Piaskowski ML, Petersen D, Dean M, Israni A, Hernandez BY, Tucker TC, Cheng I, Gonsalves L, Morris CR, Hussain SK, Lynch CF, Harris RS, Prokunina-Olsson L, Meltzer PS, Buck CB, Engels EA. 2023. Evidence for virus-mediated oncogenesis in bladder cancers arising in solid organ transplant recipients. Elife 12.

13. Ali N, Shaikh MU, Hasan S. 2011. BK Virus Associated Late Onset Haemorrhagic Cystitis After Allogeneic Peripheral Blood Stem Cell Transplant. Indian J Hematol Blood Transfus 27:177–9.

14. Sood P, Senanayake S, Sujeet K, Medipalli R, Van-Why SK, Cronin DC, Johnson CP, Hariharan S. 2013. Donor and recipient BKV-specific IgG antibody and posttransplantation BKV infection: a prospective single-center study. Transplantation 95:896–902.

15. Abend JR, Changala M, Sathe A, Casey F, Kistler A, Chandran S, Howard A, Wojciechowski D. 2017. Correlation of BK Virus Neutralizing Serostatus With the Incidence of BK Viremia in Kidney Transplant Recipients. Transplantation 101:1495–1505.

16. Hirsch HH, Randhawa PS, Practice ASTIDCo. 2019. BK polyomavirus in solid organ transplantation-Guidelines from the American Society of Transplantation Infectious Diseases Community of Practice. Clin Transplant 33:e13528.

17. Solis M, Velay A, Porcher R, Domingo-Calap P, Soulier E, Joly M, Meddeb M, Kack-Kack W, Moulin B, Bahram S, Stoll-Keller F, Barth H, Caillard S, Fafi-Kremer S. 2018. Neutralizing Antibody-Mediated Response and Risk of BK Virus-Associated Nephropathy. J Am Soc Nephrol 29:326–334.

18. Helle F, Aubry A, Morel V, Descamps V, Demey B, Brochot E. 2024. Neutralizing Antibodies Targeting BK Polyomavirus: Clinical Importance and Therapeutic Potential for Kidney Transplant Recipients. J Am Soc Nephrol 35:1425–1433.

19. D’Costa M, Vathsala A, Wong E, Chang Z, Ng A, Sran H. 2025. Intravenous Immunoglobulin as Adjunctive Treatment for BKDNAemia. American Journal of Transplantation 25:S17 (https://www.amjtransplant.org/article/S1600-6135(25)00401-0/fulltext)

20. Macdonald LE, Karow M, Stevens S, Auerbach W, Poueymirou WT, Yasenchak J, Frendewey D, Valenzuela DM, Giallourakis CC, Alt FW, Yancopoulos GD, Murphy AJ. 2014. Precise and in situ genetic humanization of 6 Mb of mouse immunoglobulin genes. Proc Natl Acad Sci U S A 111:5147–52.

21. Murphy AJ, Macdonald LE, Stevens S, Karow M, Dore AT, Pobursky K, Huang TT, Poueymirou WT, Esau L, Meola M, Mikulka W, Krueger P, Fairhurst J, Valenzuela DM, Papadopoulos N, Yancopoulos GD. 2014. Mice with megabase humanization of their immunoglobulin genes generate antibodies as efficiently as normal mice. Proc Natl Acad Sci U S A 111:5153–8.

22. Peretti A, Geoghegan EM, Pastrana DV, Smola S, Feld P, Sauter M, Lohse S, Ramesh M, Lim ES, Wang D, Borgogna C, FitzGerald PC, Bliskovsky V, Starrett GJ, Law EK, Harris RS, Killian JK, Zhu J, Pineda M, Meltzer PS, Boldorini R, Gariglio M, Buck CB. 2018. Characterization of BK Polyomaviruses from Kidney Transplant Recipients Suggests a Role for APOBEC3 in Driving In-Host Virus Evolution. Cell Host Microbe 23:628–635 e7.

23. Pastrana DV, Brennan DC, Cuburu N, Storch GA, Viscidi RP, Randhawa PS, Buck CB. 2012. Neutralization serotyping of BK polyomavirus infection in kidney transplant recipients. PLoS Pathog 8:e1002650.

24. Parsons AJ, Ophir SI, Duty JA, Kraus TA, Stein KR, Moran TM, Tortorella D. 2022. Development of broadly neutralizing antibodies targeting the cytomegalovirus subdominant antigen gH. Commun Biol 5:387.

25. Liu S, Wang S, Lu S. 2018. Using DNA Immunization to Elicit Monoclonal Antibodies in Mice, Rabbits, and Humans. Hum Gene Ther 29:997–1003.

26. Greenfield EA. 2022. Immunizing Animals. Cold Spring Harb Protoc 2022:Pdb top100180.

27. Irani V, Guy AJ, Andrew D, Beeson JG, Ramsland PA, Richards JS. 2015. Molecular properties of human IgG subclasses and their implications for designing therapeutic monoclonal antibodies against infectious diseases. Mol Immunol 67:171–82.

28. Victora GD, Nussenzweig MC. 2012. Germinal centers. Annu Rev Immunol 30:429–57.

29. Pastrana DV, Pumphrey KA, Cuburu N, Schowalter RM, Buck CB. 2010. Characterization of monoclonal antibodies specific for the Merkel cell polyomavirus capsid. Virology 405:20–5.

30. Abend JR, Sathe A, Wrobel MB, Knapp M, Xu L, Zhao L, Kim P, Desai S, Nguyen A, Leber XC, Hein A, Scharenberg M, Shaul J, Ornelas E, Wong K, Pietzonka T, Sterling LM, Hodges MR, Pertel P, Traggiai E, Patick AK, Kovacs SJ. 2024. Nonclinical and clinical characterization of MAU868, a novel human-derived monoclonal neutralizing antibody targeting BK polyomavirus VP1. Am J Transplant 24:1994–2006.

31. Shen L, Peterson S, Sedaghat AR, McMahon MA, Callender M, Zhang H, Zhou Y, Pitt E, Anderson KS, Acosta EP, Siliciano RF. 2008. Dose-response curve slope sets class-specific limits on inhibitory potential of anti-HIV drugs. Nat Med 14:762–6.

32. Akiyoshi DE, Sheoran AS, Rich CM, Richard L, Chapman-Bonofiglio S, Tzipori S. 2010. Evaluation of Fab and F(ab’)2 fragments and isotype variants of a recombinant human monoclonal antibody against Shiga toxin 2. Infect Immun 78:1376–82.

33. Casadevall A, Janda A. 2012. Immunoglobulin isotype influences affinity and specificity. Proc Natl Acad Sci U S A 109:12272–3.

34. Weber M, Schmitt S, Eicher B, Seidenberg J, Rutkauskaite J, Stockli B, Townsend C, Huynh-Do U, Schachtner T, Delbue S, Mader A, Esslinger C, Hillenbrand M. 2025. A highly potent human antibody neutralizing all serotypes of BK polyomavirus. PLoS Pathog 21:e1013122.

35. Morel V, Martin E, Francois C, Helle F, Faucher J, Mourez T, Choukroun G, Duverlie G, Castelain S, Brochot E. 2017. A Simple and Reliable Strategy for BK Virus Subtyping and Subgrouping. J Clin Microbiol 55:1177–1185.

36. Randhawa P, Pastrana DV, Zeng G, Huang Y, Shapiro R, Sood P, Puttarajappa C, Berger M, Hariharan S, Buck CB. 2015. Commercially available immunoglobulins contain virus neutralizing antibodies against all major genotypes of polyomavirus BK. Am J Transplant 15:1014–20.

37. Velay A, Solis M, Benotmane I, Gantner P, Soulier E, Moulin B, Caillard S, Fafi-Kremer S. 2019. Intravenous Immunoglobulin Administration Significantly Increases BKPyV Genotype-Specific Neutralizing Antibody Titers in Kidney Transplant Recipients. Antimicrob Agents Chemother 63.

38. Sato N, Shiraki A, Mori KP, Sakai K, Takemura Y, Yanagita M, Imoto S, Tanabe K, Shiraki K. 2024. Preemptive intravenous human immunoglobulin G suppresses BK polyomavirus replication and spread of infection in vitro. Am J Transplant 24:765–773.

39. Vu D, Shah T, Ansari J, Naraghi R, Min D. 2015. Efficacy of intravenous immunoglobulin in the treatment of persistent BK viremia and BK virus nephropathy in renal transplant recipients. Transplant Proc 47:394–8.

40. Mohammad D, Kim DY, Baracco R, Kapur G, Jain A. 2022. Treatment of BK virus with a stepwise immunosuppression reduction and intravenous immunoglobulin in pediatric kidney transplant. Pediatr Transplant 26:e14241.

41. Mosca M, Bacchetta J, Chamouard V, Rascle P, Dubois V, Paul S, Mekki Y, Picard C, Bertholet-Thomas A, Ranchin B, Sellier-Leclerc AL. 2023. IVIg therapy in the management of BK virus infections in pediatric kidney transplant patients. Arch Pediatr 30:165–171.

42. Benotmane I, Solis M, Velay A, Cognard N, Olagne J, Gautier Vargas G, Perrin P, Marx D, Soulier E, Gallais F, Moulin B, Fafi-Kremer S, Caillard S. 2021. Intravenous immunoglobulin as a preventive strategy against BK virus viremia and BKV-associated nephropathy in kidney transplant recipients-Results from a proof-of-concept study. Am J Transplant 21:329–337.

43. Karatas M, Tatar E, Okut G, Yildirim AM, Kocabas E, Tasli Alkan F, Simsek C, Dogan SM, Uslu A. 2024. Efficacy of mTOR Inhibitors and Intravenous Immunoglobulin for Treatment of Polyoma BK Nephropathy in Kidney Transplant Recipients: A Biopsy-Proven Study. Exp Clin Transplant 22:118–127.

44. Moon J, Chang Y, Shah T, Min DI. 2020. Effects of intravenous immunoglobulin therapy and Fc gamma receptor polymorphisms on BK virus nephropathy in kidney transplant recipients. Transpl Infect Dis 22:e13300.

45. Kable K, Davies CD, O’Connell P J, Chapman JR, Nankivell BJ. 2017. Clearance of BK Virus Nephropathy by Combination Antiviral Therapy With Intravenous Immunoglobulin. Transplant Direct 3:e142.

46. Rasaei N, Malekmakan L, Gholamabbas G, Abdizadeh P. 2023. Comparative Study of Intravenous Immunoglobulin and Leflunomide Combination Therapy With Intravenous Immunoglobulin Single Therapy in Kidney Transplant Patients With BK Virus Infection: Single-Center Clinical Trial. Exp Clin Transplant 21:814–819.

47. Keller N, Duquennoy S, Conrad A, Fafi-Kremer S, Morelon E, Bouvier N, Moulin B, Hurault De Ligny B, Caillard S. 2019. Clinical utility of leflunomide for BK polyomavirus associated nephropathy in kidney transplant recipients: A multicenter retrospective study. Transpl Infect Dis 21:e13058.

48. Vincent K. 2022. Vera Therapeutics Reports Positive Interim Phase 2 Data Showing MAU868 Has Significant BK Antiviral Activity in Kidney Transplant Patients, p 2, https://www.globenewswire.com/news-release/2022/06/04/2456292/0/en/Vera-Therapeutics-Reports-Positive-Interim-Phase-2-Data-Showing-MAU868-Has-Significant-BK-Antiviral-Activity-in-Kidney-Transplant-Patients.html.

