## Supplementary material for "Broadly neutralizing human monoclonal antibodies against BK polyomavirus genotypes": Text is within the figure

### Supplemental Figure 1

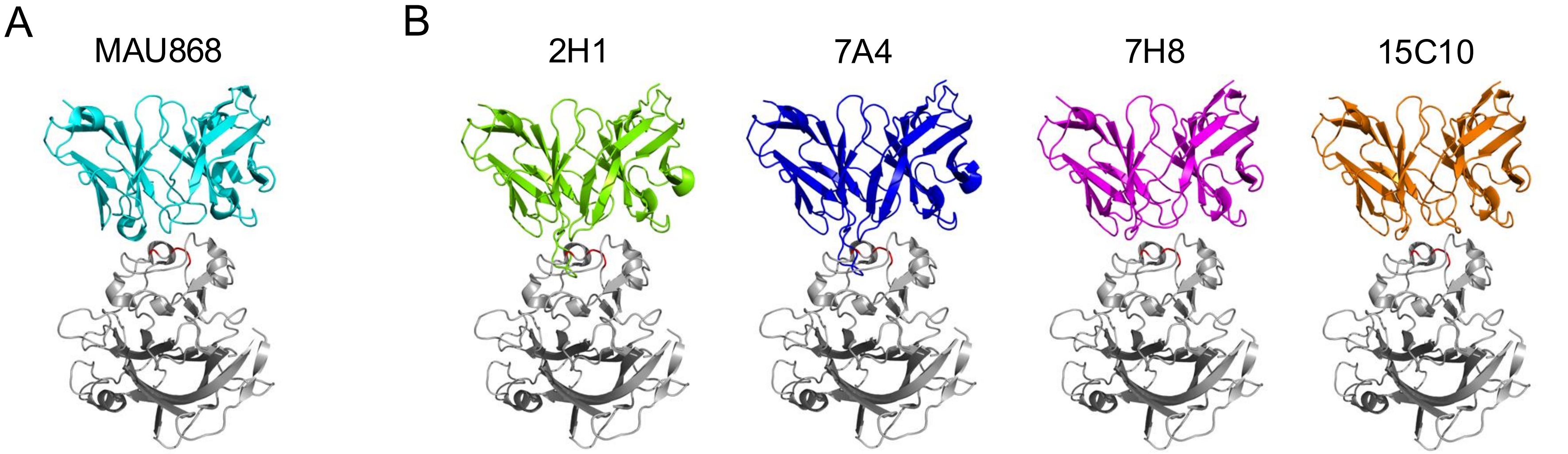


**<u>Supplemental Figure 1.</u>** <u>ColabFold predictions of mAb binding to BKV VP1 monomer via MAU868 binding pocket.</u> **A**) Representation of crystal structure of MAU868 single-chain variable fragment (colored as cyan) bound to BKV VP1 monomer (colored as grey), highlighting interactions with three key contact residues on VP1: Y169, R170, and K172 (colored as red) (30). **B**) Structural predictions of cross-reactive clones 2H1 (green), 7A4 (blue), 7H8 (magenta), and 15C10 (orange) were generated using ColabFold due to its faster model generation compared to AlphaFold. Predictive structures followed the PDB100 template and the highest ranked model of 5 relaxes was chosen for alignment. mAb predictions were then aligned to the crystal structure, resulting in effective overlap with MAU868 (RMSD range 0.322 Å to 0.651 Å). The alignment illustrates structural dissimilarity of mabs 2H1 (green), 7A4 (blue), 7H8 (magenta), and 15C10 (orange) to MAU868 in the binding region of MAU868 to the VP1 monomer. These results suggest that mAbs 2H1 (green), 7A4 (blue), 7H8 (magenta), and 15C10 (orange) likely interact to a distinct site of BKV VP1 than MAU868.
