## Supplementary material for "Broadly neutralizing human monoclonal antibodies against BK polyomavirus genotypes": N/A

### Table S1: Summary of broadly neutralizing BKV clonotype mAbs

| Clone | Heavy Chain (CDR3) Length | Heavy Chain (CDR3) Seq | Heavy D Gene | Heavy J Gene | Clonotype | Fusion |
|---|---|---|---|---|---|---|
| 2B9 | 14 | DPYNWNHGVYG MDV | IGHV3-23 | IGHJ6 | A | 1 |
| 2F11* | 10 | HARSWNYVAY | IGHV5-51 | IGHJ4 | B | |
| 3D6* | 10 | HARSWNYVAY | IGHV5-51 | IGHJ4 | B | 2 |
| 4F1 | 10 | HARNWNNVAY | IGHV5-51 | IGHJ4 | C | |
| 2G2* | 12 | DRFLEWVEGFDP | IGHV3-23 | IGHJ5 | D | |
| 14F5* | 12 | DRFLEWVEGFDP | IGHV3-23 | IGHJ5 | D | 3 |
| 15C10* | 12 | DRFLEWVEGFDP | IGHV3-23 | IGHJ5 | D | |
| 9G9 | 12 | DRFLEWVEGFDS | IGHV3-23 | IGHJ5 | D | |
| 1H10 | 11 | ERRDGHKIFDC | IGHV3-11 | IGHJ4 | E | |
| 2F7* | 11 | ERRDGHKIFDC | IGHV3-11 | IGHJ4 | E | |
| 4F7 | 11 | ERRDGHKIFDC | IGHV3-11 | IGHJ4 | E | |
| 2H1 | 11 | ERRDGHKIFDC | IGHV3-11 | IGHJ4 | E | |
| 4F9 | 11 | ERRDGHKIFDW | IGHV3-11 | IGHJ4 | E | |
| 5A8* | 11 | ERRDGHKIFDY | IGHV3-11 | IGHJ4 | E | |
| 7A4* | 11 | ERRDGHKIFDY | IGHV3-11 | IGHJ4 | E | |
| 8D10* | 11 | ERRDGHKIFDY | IGHV3-11 | IGHJ4 | E | 4 |
| 5H5 | 11 | ERRDGHKIFDW | IGHV3-11 | IGHJ4 | E | |
| 7C4 | 11 | ERRDGHKIFDW | IGHV3-11 | IGHJ4 | E | |
| 7H8 | 10 | DSSSWFSLHY | IGHV3-23 | IGHJ4 | F | |
| 8A11 | 11 | ERRSGHKIFDC | IGHV3-11 | IGHJ4 | E | |
| 8F4 | 11 | ERRDGHKIFDW | IGHV3-11 | IGHJ4 | E | |
| 8H6 | 11 | ERRDGHKIFDC | IGHV3-11 | IGHJ4 | E | |
| 9B4 | 11 | ERREGHKIFDF | IGHV3-11 | IGHJ4 | F | |

*Identical clones, respectively
